# Imaging of *Staphylococcus aureus* infections and biofilms using a selective covalent probe for the unique serine hydrolase FphE

**DOI:** 10.64898/2026.02.24.707767

**Authors:** Emily C. Woods, Tulsi Upadhyay, Ki Wan Park, Shih-Po Su, Zhen Xiao, Jianghong Rao, Tulio A. Valdez, Jeyun Jo, Matthew Bogyo

## Abstract

*Staphylococcus aureus* is the leading cause of soft tissue infections that can be treated with antibiotics. However, it can also cause significant mortality and morbidity due to systemic infections and infections of surgical implants. Implant infections typically require invasive surgery, and treatment often necessitates removal of the implant because *S. aureus* biofilms are extremely difficult to eradicate with antibiotic treatment alone. Therefore, there is a significant need for improved diagnostic tools for rapid, non-invasive confirmation of *S. aureus* infections. We recently developed an activity-based probe containing an oxadiazolone electrophile that selectively labels the *S. aureus*-specific serine hydrolase, FphE, by covalent binding to its active site serine residue. Here we describe a Cy5-labeled version of the probe, JJ-OX-012, and its characterization as an imaging agent for detecting biofilms both *in vitro* and *in vivo*. The probe labeled *S. aureus* biofilms *in vitro*, with virtually no background labeling of bacteria that lack FphE expression. Furthermore, we demonstrate that JJ-OX-012 can be used for non-invasive fluorescent imaging as a way to detect *S. aureus* biofilms *in vivo*. Overall, these findings support the potential for using covalent probes targeting FphE as imaging agents for rapid detection and diagnosis of staphylococcal infections *in vivo*.

## INTRODUCTION

*Staphylococcus aureus* is a Gram-positive bacterium that causes significant morbidity and mortality worldwide^1^. One aspect of *S. aureus* pathogenesis that contributes to its devastating health consequences is the ability to form biofilms. Due to metabolic changes and structural protection within biofilms, bacteria in biofilms have increased levels of antibiotic resistance and decreased susceptibility to host immune clearance^2,3^. Therefore, traditional treatment with antibiotics is often inadequate to clear an infection, and a surgical procedure to physically remove biofilm-associated infections may be necessary^3^.

*S. aureus* biofilms have a strong tendency to form on implanted medical devices, such as orthopedic hardware or cardiac implants ^4^. Definitive diagnosis of implant infections typically requires surgery to obtain samples from the implant for bacterial culture or PCR^5^. This process is highly invasive and slow, resulting in a significant need for diagnostic tools for non-invasive identification of surgical implant infections.

Efforts to develop imaging tools for *S. aureus* biofilms have focused on antibodies^6–8^, antibiotic-based probes^8^ or DNA-based probes^9^. Covalent small molecule probes offer advantages over these types of probes as a result of cheaper production cost, favorable pharmacokinetics, extended target occupancy, and low rates of resistance^10,11^. Although some small molecule or peptide-based probes have been successfully applied to imaging *Pseudomonas aeruginosa* biofilms^12–14^, similar tools have not previously been used for *S. aureus* biofilms.

While covalent binding probes have great advantages as imaging agents, the key to their success is selection of both an optimal protein target and optimal reactive electrophile. Enzymes such as proteases or hydrolases are ideal imaging probe targets because they possess highly nucleophilic residues in the active site that covalently react with a weak electrophile. In addition, it is possible to identify specific hydrolase enzymes that are unique to *S. aureus*, and that are expressed in biofilms. We have recently determined that the serine hydrolase, FphE meets these criteria. It is a member of the fluorophosphonate binding family of serine hydrolases that are expressed mainly in *S. aureus*^15,16^. Moreover, of this family, it is the one member that lacks homologs in virtually all other bacteria and mammals. In addition, we have recently shown that FphE is highly expressed and plays an important role in biofilm formation^17^.

Our lab previously developed an oxadiazolone-based probe that selectively labels the active site of FphE in *S. aureus*^16^. In this study we optimize an oxadiazolone probe for *in vivo* use and characterize its potential for penetrating and labeling biofilms both *in vitro* and *in vivo*. We demonstrate that this new probe, JJ-OX-012, has nanomolar potency against FphE and selectively labels *S. aureus* in both planktonic and biofilm states. In addition, compared to the prior probe, JJ-OX-012 significantly improves the signal-to-noise ratio for detecting *S. aureus* in a murine surgical site infection model. Moreover, JJ-OX-012 localizes to the site of *S. aureus* biofilms *in vivo*. The optimization of this probe for *in vivo* use resulted in an improved tool for imaging *S. aureus* biofilm infections.

## RESULTS AND DISCUSSION

### JJ-OX-012 binds covalently to FphE

Building upon our prior work that demonstrated the utility of the oxadiazolone electrophile for targeting FphE^16^, we attached Cy5 to the oxadiazolone scaffold to synthesize JJ-OX-012 (**Fig. 1A**). Because Cy5 is known to exhibit some non-specific binding^18^, we also synthesized a control compound, JJ-OX-017, which lacks the active electrophile of the oxadiazolone warhead. The synthetic methods and compound verification are available in the supplementary materials.

**Figure 1.**
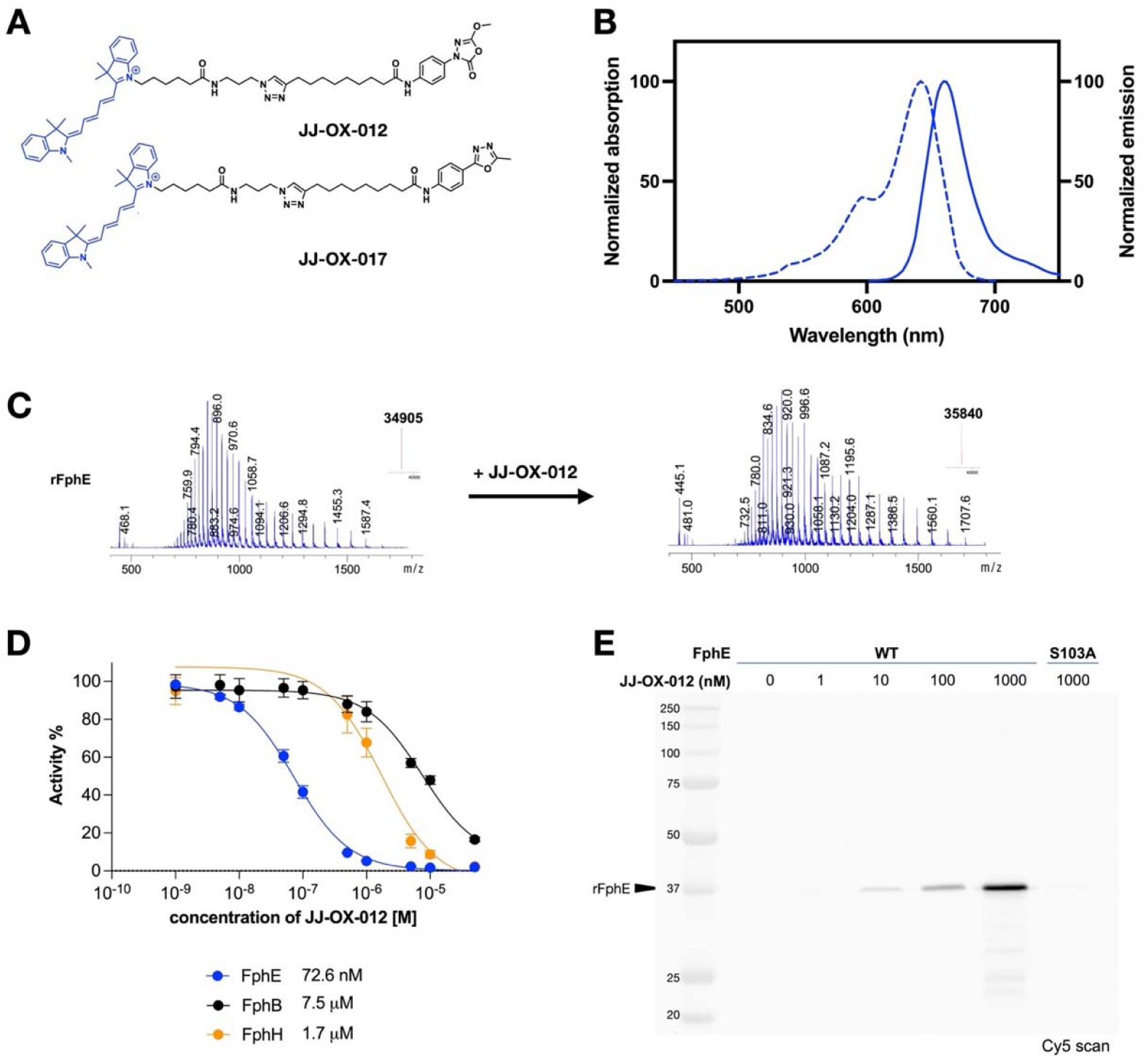
The Cy5-based probe shows selective labeling of FphE. (**a**) Chemical structures of JJ-OX-012 and JJ-OX-017. (**b**) Excitation (dotted lines) and emission (solid lines) spectra of 200 nM JJ-OX-012 as determined by fluorescence spectroscopy. (**c**) Mass spectrometry data showing the irreversible binding of JJ-OX-012 (5 µM) to rFphE (2 µM). Deconvoluted mass spectra of rFphE before (left) and after addition of compound (right) are shown. Inset in the upper right-hand corner indicates the Δm/z of 935. (**d**) IC_50_ of JJ-OX-012. Compound was incubated with FphE (blue), FphB (black), and FphH (orange) in enzyme activity assays. Percent residual activity from two independent replicates, each performed as technical triplicates, is shown as means ± S.E.M., and the calculated IC_50_ for each enzyme is listed in the legend. (**e**) Determination of active-site binding. Increasing concentrations of JJ-OX-012 were incubated with 2 µM rFphE or catalytically dead rFphE (S103A), separated on SDS-PAGE gel, and fluorescently imaged for Cy5. Ladder sizes are shown in kDa. Data is representative of three independent replicates. The corresponding Coomassie stain is shown in **Fig. S7A**.

To test whether linkage to the oxadiazolone scaffold altered the fluorescence properties of Cy5, we measured the absorbance and emission of JJ-OX-012 over a range of wavelengths (**Fig. 1B**). As expected for Cy5, we found that the excitation peak was 642 nm, and the emission peak was 660 nm.

Although previous work had established that the oxadiazolone warhead covalently binds to FphE^16^, we first tested whether addition of a Cy5 fluorophore affects binding specificity of the probe. We therefore incubated recombinant FphE (rFphE) with JJ-OX-012 and analyzed the adduct by mass spectroscopy. A mass shift of 935 Dalton was observed, which is within the range of error from the expected shift of 937 Dalton (**Fig. 1C**), indicating that JJ-OX-012 binds covalently to rFphE. In contrast, the control compound, JJ-OX-017, did not result in a mass shift of rFphE (**Fig. S1A**). Similarly, incubation of JJ-OX-012 with rFphE with the active site serine mutated to alanine (rFphE S103A) did not result in a mass shift (**Fig. S1B**). Taken together, these results confirm that JJ-OX-012 binds covalently to FphE via covalent binding of the warhead to the active-site serine.

To test the potency and specificity of JJ-OX-012, we determined the IC_50_ against recombinant FphE and the related enzymes, FphB and FphH (**Fig. 1D**). JJ-OX-012 potently inhibited rFphE activity with an IC_50_ of 72.6 nM ± 0.8 nM. This inhibition is selective, as the IC_50_ against rFphB and rFphH are two orders of magnitude higher (7.5 µM ± 3.0 µM and 1.9 µM ± 0.6 µM, respectively). In contrast, the control compound, JJ-OX-017, did not inhibit rFphE (IC_50_ > 50 µM; **Fig. S2**), further confirming the critical role of covalent binding between the electrophile of JJ-OX-012 with FphE in this interaction.

As an initial assessment of the ability to image FphE with JJ-OX-012, we incubated varying concentrations of JJ-OX-012 with rFphE and visualized the Cy5-labeled proteins after resolution by SDS-PAGE (**Fig. 1E**). The probe retained detectable labeling of rFphE to as low as 10 nM. In further support of the interaction between the probe and active site of the enzyme, rFphE S103A had no detectable labeling by JJ-OX-012 at the highest concentration tested (1 µM). Similarly, JJ-OX-017 did not label rFphE up to 1 µM (**Fig. S3**), indicating that the electrophile is necessary for binding to the enzyme.

### JJ-OX-012 binds selectively to FphE in *S. aureus*

Having established that JJ-OX-012 covalently binds to rFphE at nanomolar concentrations, we next sought to determine if it is also able to label FphE expressed in live *S. aureus* cells. Incubation of various concentrations of the probe with *S. aureus* wild-type (WT) bacteria resulted in strong labeling of a ∼31 kDa protein (**Fig. 2A**). This protein is absent in the *S. aureus fphE*::Tn mutant strain, confirming its identity as FphE. At the highest concentrations tested (100 nM), as well as in the *fphE* mutant, the faintly labeled protein at ∼38 kDa likely corresponds to the related enzyme, FphB^17^. Thus, although FphE is the primary target of JJ-OX-012, this probe can react with FphB when FphE is absent or when the probe is used at high concentrations. Incubation of the bacteria with the negative control probe JJ-OX-017 did not result in any labeling, further confirming the fluorescent signal in labeled bacteria results from covalent modification of FphE by JJ-OX-012.

**Figure 2.**
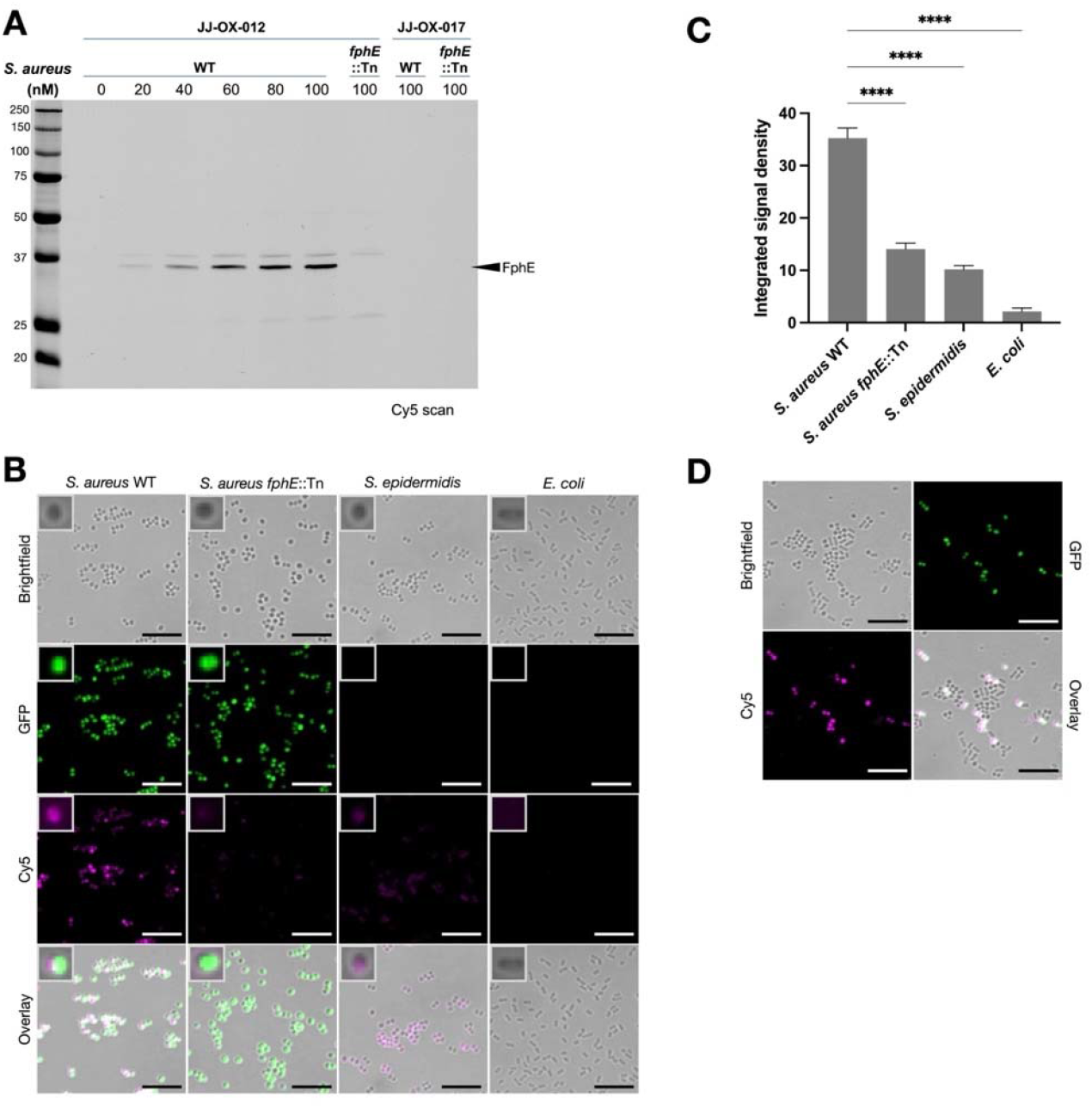
JJ-OX-012 binds selectively to FphE in *S. aureus* planktonic cells. (**a**) JJ-OX-012 labeling of live *S. aureus. S. aureus* USA300 wild-type (WT) or *fphE* transposon mutant (*fphE*::Tn) was incubated with 0 to 100 nM of JJ-OX-012, washed, lysed, and equal amounts of total protein were separated on SDS-PAGE gel and fluorescently imaged for Cy5. Ladder sizes are shown in kDa. Data is representative of three independent replicates. The corresponding Coomassie stain is shown in **Fig. S7C** (**b**) Planktonic cell fluorescent microscopy. Stationary phase cultures of *S. aureus* USA300 expressing GFP, *S. aureus fphE*::Tn expressing GFP, *S. epidermidis*, and *E. coli* were incubated with 50 nM JJ-OX-012 for 2 hours, washed, and imaged by confocal microscopy. Brightfield, GFP, Cy5, and BF/Cy5/GFP composite images are shown. Scale bars represent 10 µm. Inset in upper left-hand corner shows single cells for morphology. Images are representative of three independent replicates. (**c**) Mean Cy5 signal intensity within bacteria (n = 100 per strain/species). Results were analyzed by one-way ANOVA with Dunnett’s multiple comparisons test. **** indicates *p* < 0.0001. (**d**) Fluorescent microscopy of mixed bacteria. Equal concentrations (based on OD_600_) of *S. aureus* USA300 expressing GFP, *S. epidermidis*, and *E. coli* were mixed and labeled as described for (**b**). Brightfield, GFP, Cy5, and BF/Cy5/GFP composite images are shown. Scale bars represent 10 µm. Images are representative of three independent replicates.

Next, we performed fluorescent microscopy on bacteria incubated with JJ-OX-012. Similar to the results observed in the gel experiments, *S. aureus* WT bacteria were strongly labeled by the probe, whereas the *fphE*::Tn mutant was only weakly labeled (**Fig. 2B**). We also used the probe to label *Staphylococcus epidermidis*, a related staphylococcal species that lacks an FphE homolog^16^ and found that this bacterial species also had only weak labeling. The weak labeling of the *fphE* mutant and *S. epidermidis* suggests some binding to related targets, such as FphB, when FphE is absent, consistent with our biochemical analysis of probe labeling using the gel readout. Lastly, to validate the specificity of the probe, we tested the Gram-negative bacterium, *Escherichia coli*, which lacks Fph homologs^16^. *E. coli* did not show any detectable labeling by JJ-OX-012 (**Fig. 2C)**. Incubation with JJ-OX-017 did not result in significant labeling of *S. aureus, S. epidermidis*, or *E. coli* (**Fig. S4**). Moreover, the probe specifically labeled *S. aureus* in a mixture of *S. aureus* WT expressing GFP, *S. epidermidis*, and *E. coli* were labeled with JJ-OX-012 (**Fig. 2D**). Together, these results demonstrate the selectivity of JJ-OX-012 for *S. aureus* FphE.

### JJ-OX-012 labels *S. aureus* biofilms *in vitro*

Biofilms are complex 3D structures, containing polysaccharides, proteins, lipids, and extracellular DNA. Due to this unique composition, alterations in parameters like hydrophobicity and pH make them notoriously difficult for compounds to penetrate^19^. We therefore wanted to test whether JJ-OX-012 can label *S. aureus* when grown as a biofilm, and whether the probe retains its selectivity for *S. aureus* in biofilms. JJ-OX-012 retained robust labeling of *S. aureus* WT in biofilms (**Fig. 3A**).

**Figure 3.**
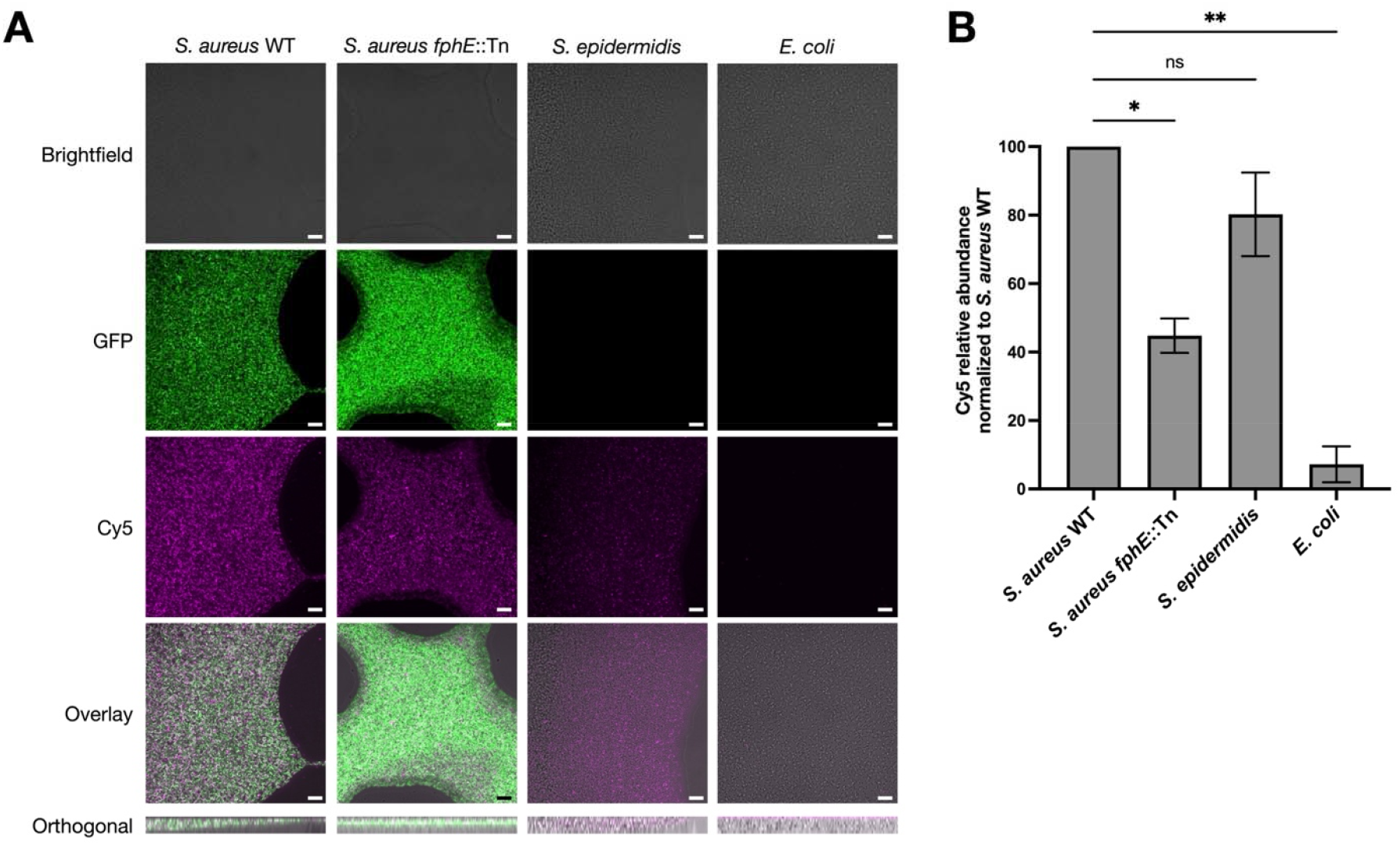
JJ-OX-012 labels *S. aureus* biofilms *in vitro*. (**a**) *S. aureus* USA300 expressing GFP, *S. aureus* fphE::Tn expressing GFP, *S. epidermidis*, and *E. coli* were grown as biofilms on glass-bottomed cell culture dishes, washed to remove planktonic cells, incubated with 25 nM JJ-OX-012 or JJ-OX-017 for 2 hours, washed, and imaged by confocal microscopy. Brightfield (BF), GFP, Cy5, and BF/Cy5 or GFP/Cy5 composite (overlay) images are shown as applicable. A representative orthogonal slice demonstrates biofilm depth. Scale bars represent 10 µm. Images are representative of three independent replicates. Crystal violet staining as further validation of biofilm formation is shown in **Fig. S5** (**b**) Quantification of Cy5 signal in biofilms. Relative abundance indicates the percentage of total biomass, as detected by brightfield, positive for Cy5 labeling. Values are normalized to *S. aureus* WT and shown as means ± S.E.M. from three independent replicates. Results were analyzed by one-way ANOVA with Dunnett’s post-test. * indicates *p* < 0.05, ** indicates *p* < 0.01, ns = not significant.

These results indicate that the probe can access FphE even within the biofilm environment. However, both the *S. aureus fphE*::Tn mutant and *S. epidermidis* were also labeled by JJ-OX-012, albeit to a lesser extent (45% and 80% of WT labeling, respectively; **Fig. 3B**). The labeling in these two strains is likely due to binding to FphB, which we observed as a secondary target in the *fphE*::Tn mutant in gel labeling (**Fig. 2A**). Importantly, biofilms of *E. coli*, which lacks homologs to any of the Fph proteins^16^, were minimally labeled (7% of WT labeling; **Fig. 3B**). The control compound JJ-OX-017 also labeled *S. aureus* WT, *S. aureus fphE*::Tn, and *S. epidermidis* but had minimal labeling of *E. coli* (**Fig. S6A**), and none of the strains had significantly different binding compared to WT (**Fig. S6B**). Because JJ-OX-017 does not label planktonic cells (**Fig. 2A and S3**), this result suggests that the dye-labeled molecules may get retained in the biofilm due to non-specific binding interactions. The differences in labeling across strains seen with JJ-OX-012 (**Fig. 3B**) which are absent with JJ-OX-017 labeling (**Fig. S6B**) suggest that, despite some non-specific binding of the dye in biofilms, the oxadiazolone warhead enables selective engagement with the FphE target.

### JJ-OX-012 localizes to the site of *S. aureus* infection in a mouse implant infection model^16^

Using a mouse surgical site infection model, we tested whether JJ-OX-012 can selectively detect *S. aureus in vivo*. In this model, an incision is made over each thigh, and bacteria are injected subcutaneously before the incision is sutured, with *S. aureus* injected on the left side, and *E. coli* injected on the right side. After systemic injection of the probe, Cy5 signal exclusively localized to the site of *S. aureus* infection (**Fig. 4A-B**). These results indicate that the probe retains selectivity for *S. aureus in vivo* and does not merely localize to areas of inflammation. Moreover, the ratio of the signal from the *S. aureus* infected thigh over the signal from the *E. coli* infected thigh was 6.9, representing a 2.5-fold improvement to the signal-to-noise ratio compared to the prior TAMRA-based version of this probe^16^. Thus, the use of the Cy5 fluorophore on the oxadiazolone scaffold significantly improved the imaging properties of this probe for *in vivo* applications.

**Figure 4.**
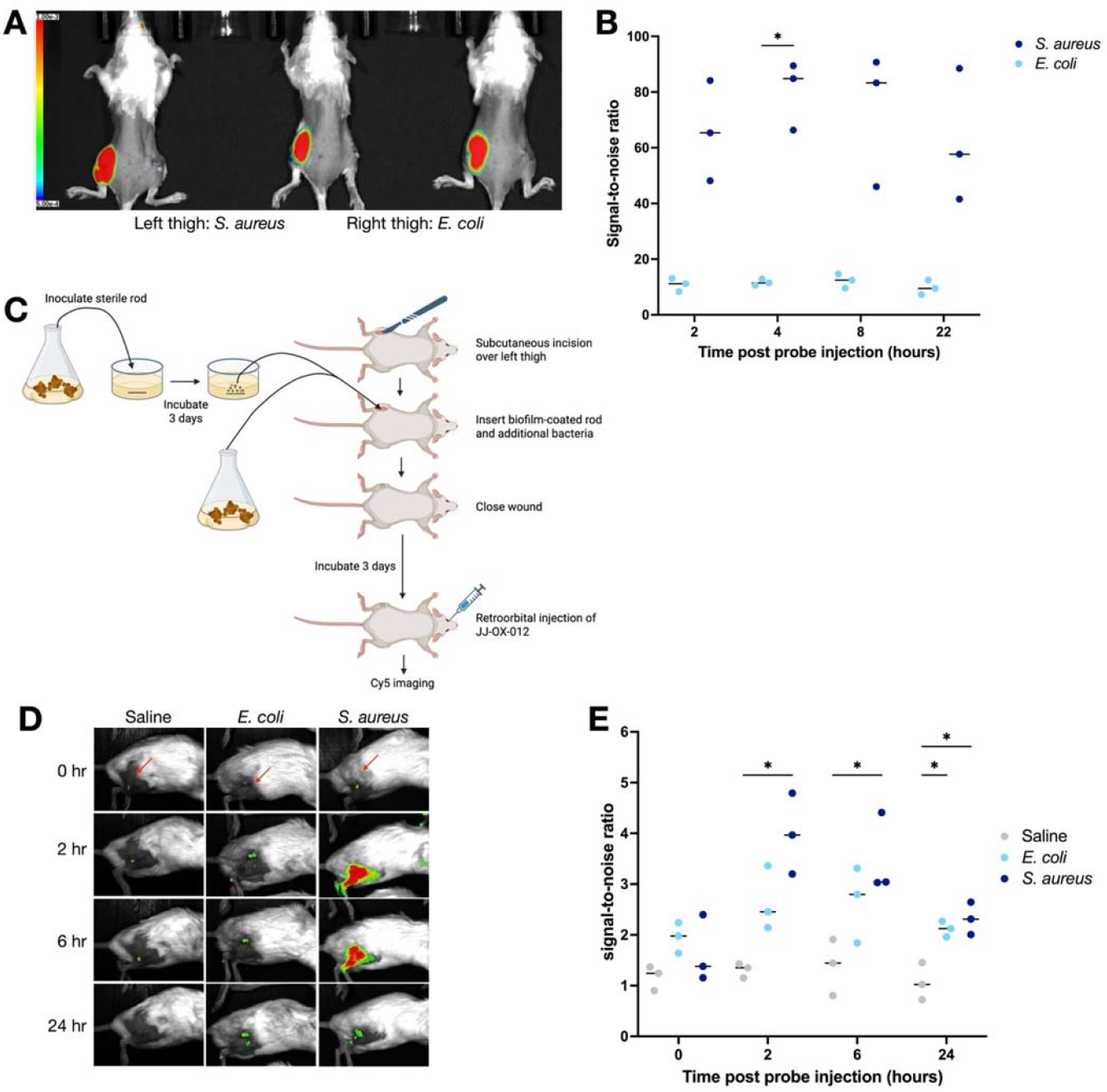
JJ-OX-012 localizes to the site of *S. aureus* infection in a surgical site infection model and implant biofilm infection model. (**a**) Cutaneous incisions were made over both thighs of BALB/c mice (n = 3) and *S. aureus* (left side) or *E. coli* (right side) was injected subcutaneously to simulate a surgical site infection. 10 nmol JJ-OX-012 was injected venously and mice were imaged serially for Cy5 signal. Cy5 imaging at 2 hours is shown with heat map scale for total efficiency displayed on the left. (**b**) Signal-to-noise ratio was calculated as the fluorescent efficiency over the region of interest (left thigh for *S. aureus* and right thigh for *E. coli*) divided by the fluorescent efficiency over an uninvolved region on the back of the same mouse. Results were analyzed by two-way ANOVA with Sidak’s multiple comparison test. * indicates *p* < 0.05. (**c**) Schematic of surgical implant model. Biofilms of *S. aureus* or *E. coli* were pre-grown on sterile titanium rods and inserted subcutaneously over the thighs of BALB/c mice (n = 3 per condition) along with additional bacteria or sterile saline and allowed to incubate for 3 days. 10 nmol of JJ-OX-012 was injected venously and mice imaged serially for Cy5 signal. Made using BioRender. (**d**) A representative mouse for each condition of the surgical implant model is shown for all time points. Red arrows at the 0 hour time point indicate the location of the rod based on corresponding CT imaging (**Fig. S8**). (**e**) Results from the biofilm implant model were analyzed as described for (**b**).

To assess whether our probe could be used for *in vivo* detection of biofilms, we tested the probe in a modified mouse surgical implant infection model^20^. In this model, biofilms were grown on a titanium rod prior to subcutaneous implantation in mice. Additional bacteria were inoculated with the rods to ensure robust biofilm generation. After 3 days to allow time for biofilms to stabilize and surgical site healing, the probe was injected venously, and mice were imaged for Cy5 signal (**Fig. 4C**).

Compared to an uninfected implant, implants infected with *S. aureus* had approximately 12-fold higher signal at 2 hours post-injection (**Fig. 4D-E**), indicating that the probe selectively localizes to *S. aureus*, not just to areas of post-surgical inflammation. Similarly, compared to *E. coli-*infected implants, *S. aureus-*infected implants had 2.5-fold higher signal at 2 hours post-injection, further demonstrating the selectivity of this probe. These results suggest that JJ-OX-012 selectively binds to *S. aureus* in an *in vivo* biofilm model.

## CONCLUSIONS

*S. aureus* biofilms commonly cause surgical implant-associated infections, are challenging to treat, and typically involve invasive surgery to diagnosis^4,5^. Here we report the development of a fluorescent probe, JJ-OX-012, that has the potential to serve as a novel tool for non-invasive diagnosis of *S. aureus* biofilm infections *in vivo*. We demonstrate that this Cy5-based probe retains high selectivity and potency for the *S. aureus*-specific enzyme, FphE. This probe can selectively label *S. aureus* cells in both planktonic and biofilm forms *in vitro*. Moreover, systemic injection of JJ-OX-012 resulted in favorable signal-to-noise ratios (SNR) for selectively detecting *S. aureus* in two infection models, highlighting the advantage of the Cy5 fluorophore for *in vivo* applications. Overall, JJ-OX-012 adds a new covalent tool for non-invasive imaging of *S. aureus* biofilms, with the potential for clinical translation.

## MATERIALS AND METHODS

### Fluorescence spectroscopy

To measure the fluorescence spectra, JJ-OX-012 was diluted in 1X PBS to a final concentration of 200 nM and transferred to a quartz fluorescence cuvette (Hellma^®^, Sigma-Aldrich, Cat. No. Z600768). Excitation and emission spectra were recorded using a wavelength-calibrated FluoroMax-3 spectrofluorometer (Horiba Jobin Yvon, Edison, NJ) equipped with a 150-W ozone-free xenon arc lamp.

17

### Gel labeling analysis of purified recombinant proteins

Purified WT (2.0 µM) and S103A rFphE (2.0 µM) in 20 µL of 1X PBS solution were incubated with a range of concentrations from 0 to 1000 nM of each chemical probe for 1 h at 37 ºC. After incubation, 20 µL of 4X SDS loading buffer was added to each sample, followed by boiling at 100 ºC for 5 min. The samples were then subjected to SDS-PAGE (12%) at 120 V in an electrophoresis chamber under the ambient temperature. Fluorescence signals were detected using a GE Typhoon FLA 9000 (GE Healthcare, Pittsburgh, PA). After imaging, the gels were stained with Coomassie Brilliant Blue for total protein visualization.

### Inhibition assay

Enzyme activity was determined using fluorogenic substrate assays as previously described^17^. In brief, recombinant FphE, FphB, and FphH were expressed and purified as previously described^17,21^. JJ-OX-012 or JJ-OX-017 were mixed in concentrations ranging from 1 nM to 50 µM with rFphE (0.5 nM), rFphB (100 nM), or rFphH (5 nM) in 0.02% TritonX-100 in 1x PBS and incubated for one hour at room temperature prior to addition of 50 µM 4-methylumbelliferone-conjugated substrates. Fluorescence (λex□ =□365□nm, λem□= □455□nm) was measured every 1 minute for 60 minutes on a Cytation 3 imaging reader (BioTek, Winooski, VT, USA). To calculate IC_50_, turnover rates were determined from the linear phase of the reaction, background rate of substrate hydrolysis subtracted, and values fitted to a non-linear curve.

### Bacterial growth

Bacterial strains used in this study are listed in **Table S1**. Bacteria were grown from frozen glycerol stocks on tryptic soy agar (TSA) plates by overnight incubation at 37°C. Starter cultures were generated by transferring a single colony to tryptic soy broth (TSB; Sigma-Aldrich, St. Louis, MO) and grown shaking at 37°C. *S. aureus* strains expressing GFP from plasmid pCM29 were grown in media supplemented with 10 µg/ml chloramphenicol to maintain the plasmid.

### SDS-PAGE analysis of live *S. aureus* labeling

Wild-type (WT) and *fphE* transposon mutant (*fphE*::Tn) *Staphylococcus* aureus USA300 strains were streaked on tryptic soy agar (TSA) plates by overnight incubation at 37 °C. Three colonies from each strain were inoculated into 10 mL tryptic soy broth (TSB) and cultured to stationary phase with shaking at 37 °C. Bacterial cultures were centrifuged at 5,000 g for 10 min, and the supernatant was discarded. Bacterial pellets were washed twice in 1X PBS and resuspended in the same buffer (5 mL). Aliquots of 400 µL were prepared from each sample. Cells were incubated 0 to 100 nM of either JJ-OX-012 or JJ-OX-017 for 2 h at 37 ºC. After incubation, cells were washed twice with 1X PBS and resuspended in the same buffer. Cell lysis was performed by bead-beating at 4 ºC. The resulting lysates were added by 4X SDS loading buffer and boiled at 100 ºC for 5 min. The denatured samples were then cooled to ambient temperature and analyzed by SDS-PAGE (12%) running at 120 V in an electrophoresis chamber under ambient conditions. Protein concentrations for each sample were quantified by using the BCA assay kit (Pierce/Thermo Scientific, Rockford, IL), and equal amounts of protein were loaded into each lane. Fluorescence signals were visualized using a GE Typhoon FLA 9000 imaging system, followed by Coomassie Brilliant Blue staining for total protein detection.

### Planktonic cell labeling and fluorescent microscopy

5 mL starter cultures as described above were grown overnight, pelleted for 10 min at 1500 *g*, and washed once in 10 ml sterile 1x phosphate buffered saline (PBS, pH 7.4). The resulting pellets were resuspended in 2 ml PBS. Cultures were adjusted to the same density as each other based on OD_600_ and each strain was aliquoted into two 0.5 mL aliquots. Additionally, a mixture of equal volumes of *S. aureus* WT, *S. epidermidis*, and *E. coli* was made in duplicate. 50 nM JJ-OX-012 or JJ-OX-017 was added to each aliquot and samples incubated for 2 hours at 37°C with shaking at 300 rpm on a thermomixer in the dark. Samples were subsequently washed twice in 1 mL PBS to remove unbound probe. Samples were resuspended in 1 mL PBS and 2 µL was plated on a 1% agarose pad on top of a glass slide and covered with a cover slip. Cy5, GFP, and brightfield (via transmitted photomultiplier tube [T-PMT]) channels were captured by confocal microscopy (Stellaris 5 microscope, Leica, Deerfield, IL). Both Cy5 (excitation 649 nm) and GFP (excitation 489 nm) imaging were performed with 1.5% laser power and line accumulation = 4. Adjustments to image brightness and contrast were made using FIJI software^22^ (ImageJ2, version 2.16.0) and applied to all images equally. For signal quantification in FIJI, regions of interest (ROI) were selected based on bacterial location in the brightfield channel, and the integrated density (total fluorescent signal within ROI / area of ROI) was measured in the Cy5 channel.

### Biofilm labeling and fluorescent microscopy

5 mL starter cultures as described above were grown for approximately 8 hours. Cultures were diluted 1:10 into 150 µl of TSB supplemented with 0.5-1% glucose (1% for *S. aureus*, 0.5% for *S. epidermidis* and *E. coli*) and transferred to a 4-chamber 35 mm glass-bottomed culture dish (Cellvis, Mountain View, CA). All cultures were plated in triplicate. Cultures were incubated stationary for 40 hours at 37°C to allow biofilm formation. Supernatant was pipetted off, and the residual biofilms were gently washed twice with 1 mL sterile 1x PBS to remove non-adherent cells. To one dish, 100 µL 0.1% crystal violet was added and incubated for 30 min at 37°C. To the other two dishes, each biofilm was covered with 0.5 mL PBS with 25 nM JJ-OX-012 or JJ-OX-017 and incubated for 2 hours at 37°C in the dark.

After incubation, supernatant was pipetted off and biofilms were gently washed twice with 1 mL PBS to remove unbound probe. Crystal violet-stained dishes were imaged by standard photography over a light box. Probe-stained dishes were imaged by confocal microscopy (Zeiss Axio Observer Z1 LSM 700, Carl Zeiss GmbH, Jena, Germany). Cy5 imaging was performed with excitation 649 nm, laser power 10%, gain 600, and GFP imaging was performed with excitation 488 nm, 1% laser power, and gain 400. Brightfield images were obtained using T-PMT in the Cy5 channel. Adjustments to image brightness and contrast were made using FIJI software and applied to all images equally. Signals were analyzed in MATLAB (version R2024b; The MathWorks, Inc., Natick, MA) using the BiofilmQ add-on^23^.Total biomass was determined in the brightfield channel, and relative abundance of Cy5 labeling was determined by assessing the percent overlap of the total biomass with the biomass with Cy5 signal. The resulting relative abundance values were normalized to the *S. aureus* WT value for each experiment and converted to percentages.

### Mouse surgical site infection model

All animal models were approved by the Stanford Institutional Animal Care and Use Committee. The surgical site infection model was performed as previously described^16^. In brief, *S. aureus* USA300 and *E. coli* BW25113 were grown in TSB media to exponential growth phase (OD_600_ = 0.5), washed three times with 1X PBS, and suspended to a concentration of 5 × 10^9^ CFU/mL. Under isoflurane anesthesia, small incisions were made in the skin overlying both rear thighs of 6 - 8 week-old female BALB/c mice. For an inoculum of 5 × 10^7^ CFU, 10 µl of either the *S. aureus* culture (left thigh) or *E. coli* culture (right thigh) was injected subcutaneously at the site of incision. The incisions were then closed with interrupted 5-0 Prolene sutures. Two and a half hours after inoculation, 10 nanomoles (100 µl of a 100 µM solution) of JJ-OX-012 (prepared in 30% PEG-400 and PBS) was injected venously via the retroorbital route. Mice were imaged under anesthesia at 2, 4, 8, and 22 hours after probe injection using an IVIS (LagoX from Spectral Instruments, Tucson, AZ) with excitation at 610 nm, medium binning, long pass filter at 710 nm, and 120 second exposure time. Fluorescence signals were measured from the images as total efficiency using Aura software (Spectral Instruments), where total efficiency equals mean signal efficiency (ratio of detected emission radiance over LED excitation radiance) in the region of interest (ROI) x area of ROI. Signals were measured over each of the infected thighs and divided by the signal from an uninfected region of the back of the same animal (background signal) to determine the signal-to-noise ratio.

### Mouse subcutaneous implant infection model

Cultures of *S. aureus* USA300 and *E. coli* BW25113 were grown in TSB media until mid-log phase, then diluted 1:10 into fresh TSB with either 0.5% (for *E. coli*) or 1% (for *S. aureus*) glucose to encourage biofilm formation. Approximately 3 mm sterile titanium rods were placed in a 12-well culture plate, covered with 1 ml of the prepared bacterial cultures, and allowed to incubate for 72 hours to allow for biofilm formation.

Incisions were made in the skin overlying the left thigh of 6-week-old female BALB/c mice under isoflurane anesthesia. The pre-infected titanium rods were inserted through the incision below the skin and an additional inoculum of 10^6^ CFU *E. coli* or 10^3^ CFU *S. aureus* in 10 µl was added to promote additional biofilm formation after insertion. Incisions were closed with interrupted 5-0 Prolene sutures. Three days after rod insertion, mice were injected with 10 nanomoles (20 µl of a 500 µM solution) of JJ-OX-012 (prepared in 30% PEG-400 and PBS) via the retroorbital route. Images were taken using an IVIS at 0, 2, 6, and 24 hours after probe injection as described above. Analysis of fluorescent signals was performed as described above.

### Statistical analysis

All statistical analyses were performed using GraphPad Prism version 10.6 (GraphPad Software, San Diego, CA). Results were analyzed by one-way ANOVA with Dunnett’s post-test or two-way ANOVA with either Tukey or Sidak post-test as described in the results.

## ACKNOWLEDGEMENTS

We would like to thank all the members of the Bogyo Lab who contributed useful feedback on this work. Additional thanks to Benyam Kinde, Satinder Kaur, Beth Mills, and Antoine Dufour for their helpful discussions. This work was funded by National Institutes of Health grant R01 EB037592 (to M.B.) and R01 DC021326 (to T.A.V.). The content of this manuscript is solely the responsibility of the authors and does not reflect the official views of the National Institutes of Health.

