## Supporting Information for "Imaging of *Staphylococcus aureus* infections and biofilms using a selective covalent probe for the unique serine hydrolase FphE"

Contents:

1. Table S1. Bacterial strains used in this study.
2. Figure S1. Mass spectrometry of rFphE with JJ-OX-017 and rFphE (S103A) with JJ-OX-012.
3. Figure S2. $IC_{50}$ of JJ-OX-017.
4. Figure S3. Gel labeling of rFphE with JJ-OX-017.
5. Figure S4. Labeling of planktonic cells by JJ-OX-017.
6. Figure S5. Crystal violet staining of biofilms.
7. Figure S6. Labeling of biofilms by JJ-OX-017.
8. Figure S7. Coomassie stains of SDS-PAGE gels.
9. Figure S8. CT scan of mice with titanium rods.
10. Chemical synthesis
    a) General methods
    b) Synthesis of JJ-OX-012
    c) Synthesis of JJ-OX-16 and JJ-OX-017
    d) LC traces of JJ-OX-012 and JJ-OX-017
    e) NMR spectra
11. References

**Table S1. Bacterial strains used in this study**

| **Strain** | **Description** | **Reference/Source** |
|---|---|---|
| *S. aureus* USA 300 LAC | Wild-type USA300 Los Angeles County (LAC) clone; cured of antibiotic resistance plasmid; MRSA | Manuel Amieva, Stanford University [1] |
| *S. aureus* USA 300 *fphE*::Tn | Transposon insertion mutant in *SAUSA300_2518*; $Ery^R$, $Linc^R$; MRSA | Nebraska Transposon Mutant Library |
| *S. aureus* USA 300 LAC + pCM29 | *S. aureus* USA 300 LAC with plasmid expressing GFP under control of the constitutive *sarAP1* promoter; $Cm^R$; MRSA | Manuel Amieva, Stanford University [2] |
| *S. aureus* USA 300 *fphE*::Tn + pCM29 | Transposon insertion mutant in *SAUSA300_2518* with plasmid expressing GFP under control of the constitutive sarAP1 promoter; $Ery^R$, $Linc^R$, $Cm^R$; MRSA | This study |
| *Staphylococcus epidermidis* RP62A (ATCC 35984) | Wild-type strain, biofilm-forming | Michael Fischbach, Stanford University |
| *Escherichia coli* ATCC 25922 | Wild-type strain, biofilm-forming | American Type Culture Collection |

Abbreviations: MRSA - methicillin resistant *S.* aureus, MSSA - methicillin sensitive *S.* aureus, $Ery^R$ - erythromycin resistant, $Linc^R$ - lincosamide resistant, $Cm^R$ - chloramphenicol resistant

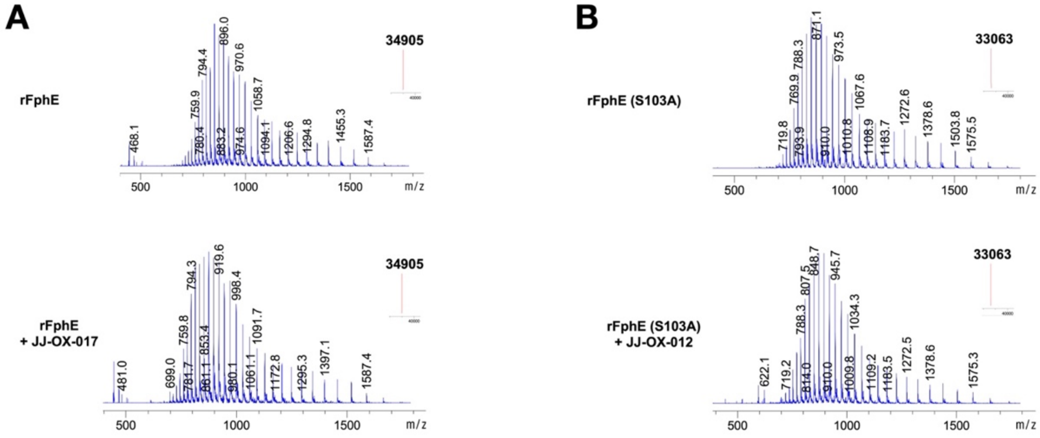


**Figure S1. Mass spectrometry of rFphE with JJ-OX-017 and rFphE (S103A) with JJ-OX-012.** (**a**) Recombinant FphE (rFphE) 2 µM was incubated with 5 µM JJ-OX-017 for 1 hour at RT and deconvoluted mass spectra of rFphE before (top) and after addition of compound (bottom) were obtained. Inset in the upper right-hand corner indicates the expected Δm/z of 0. (**b**) Recombinant FphE with active site mutation (rFphE (S103A)) 2 µM was incubated with 5 µM JJ-OX-012 for 1 hour at RT and deconvoluted mass spectra of the enzyme before (top) and after addition of compound (bottom) were obtained. Inset in the upper right-hand corner indicates the expected Δm/z of 0.

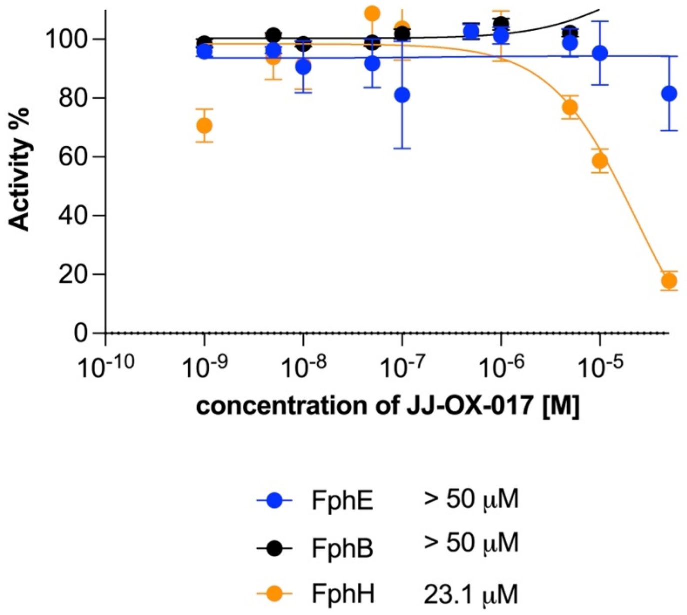


**Figure S2. $IC_{50}$ of JJ-OX-017.** Compound was incubated with FphE (blue), FphB (black), and FphH (orange) in enzyme activity assays. Percent residual activity from three replicates are shown as means ± S.E.M., and the calculated $IC_{50}$ for each enzyme is listed in the legend.

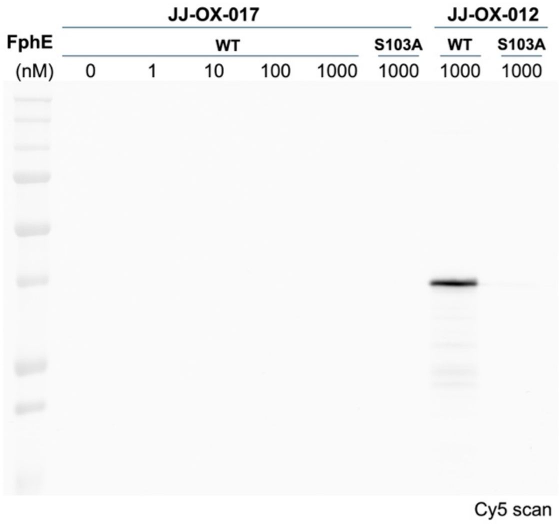


**Figure S3. Gel labeling of rFphE with JJ-OX-017.** Concentrations of JJ-OX-017 ranging from 0 to 1000 nM were incubated with 2 μM rFphE or catalytically dead rFphE (S103A), separated on SDS-PAGE gel, and fluorescently imaged for Cy5. Labeling with 1000 nM JJ-OX-012 is included for comparison. Data is representative of three independent replicates. The corresponding Coomassie stain is available in **Fig. S7B**.

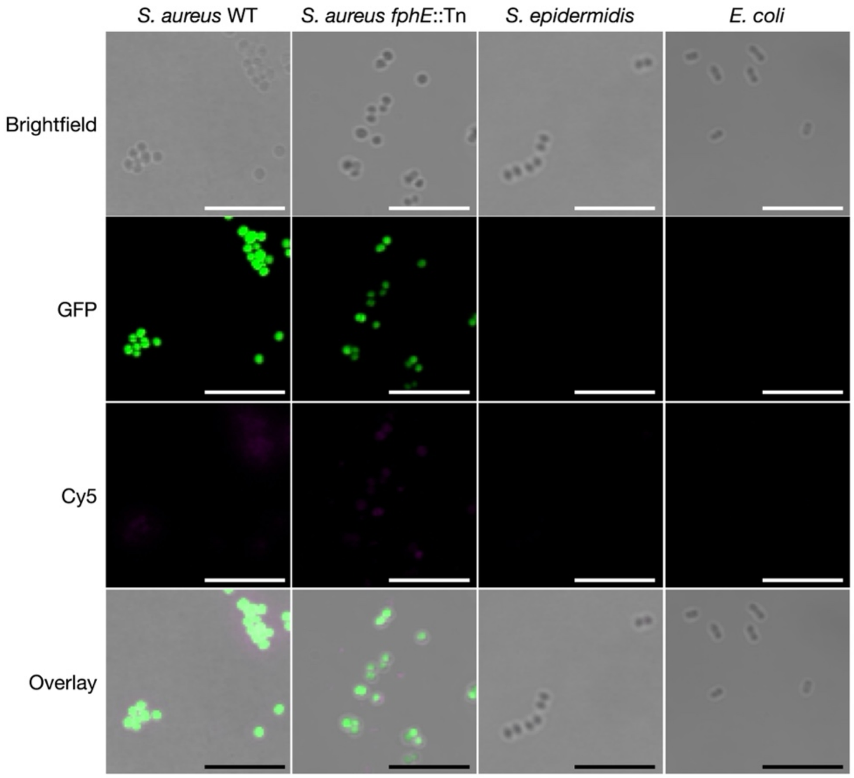


**Figure S4. Labeling of planktonic cells by JJ-OX-017.** Planktonic cell fluorescent microscopy. Stationary phase cultures of *S. aureus* USA300 expressing GFP, *S. aureus fphE*::Tn expressing GFP, *S. epidermidis*, and *E. coli* were incubated with 50 nM JJ-OX-017 for 2 hours, washed, and imaged by confocal microscopy. Brightfield, GFP, Cy5, and BF/Cy5/GFP composite images are shown. Scale bars represent 10 µm. Images are representative of three independent replicates.

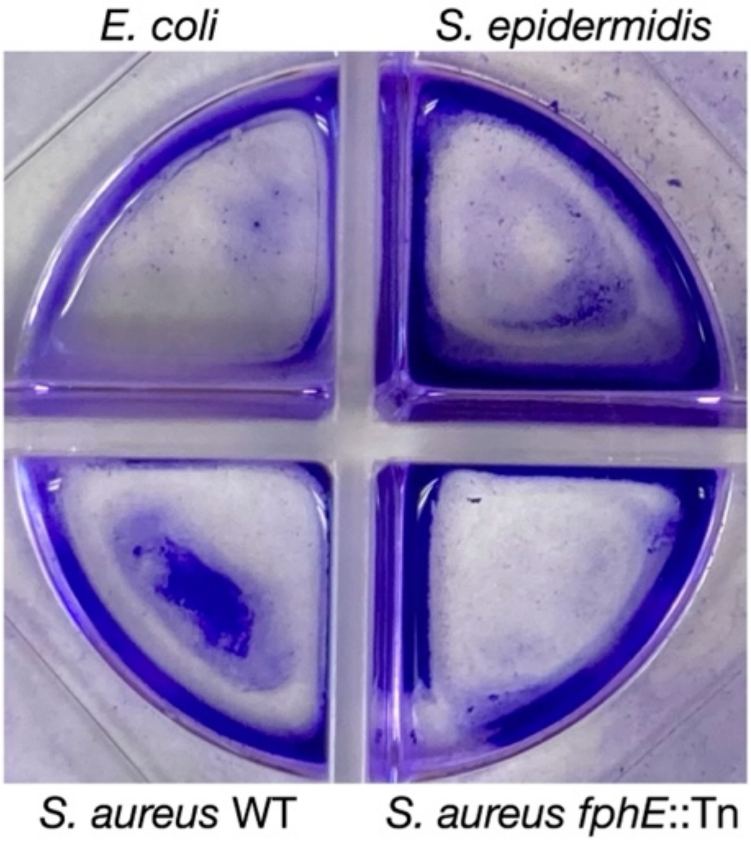


**Figure S5. Crystal violet staining of biofilms.** As a control to verify biofilm formation, cultures for confocal microscopy were grown in duplicate in four-chamber glass-bottom microscopy dishes. One set was used for confocal microscopy and the other set (shown here) was used for crystal violet staining. Biofilms were washed once with PBS to remove planktonic cells, incubated with 0.1% crystal violet for 30 min at 37˚C, washed twice with PBS, and photographed.

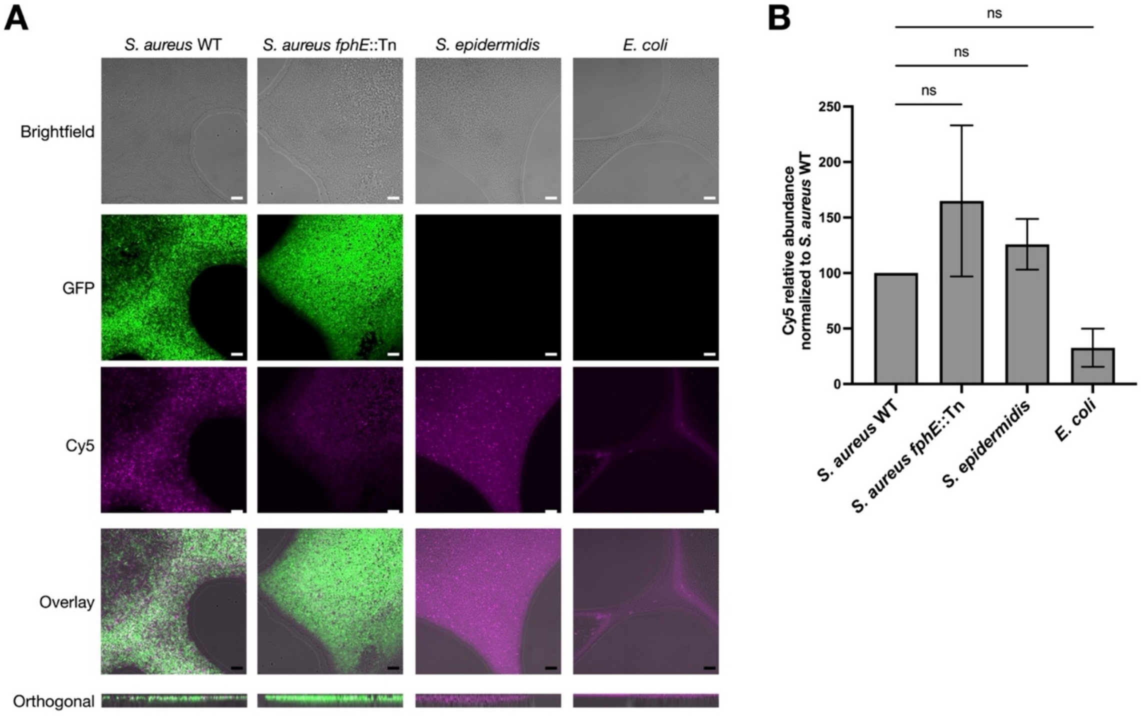


**Figure S6. Labeling of biofilms by JJ-OX-017.** (**a**) *S. aureus* USA300 expressing GFP, *S. aureus* fphE::Tn expressing GFP, *S. epidermidis*, and *E. coli* were grown as biofilms on glass-bottomed cell culture dishes, washed to remove planktonic cells, incubated with 25 nM JJ-OX-012 or JJ-OX-017 for 2 hours, washed, and imaged by confocal microscopy. Brightfield (BF), GFP, Cy5, and BF/Cy5 or GFP/Cy5 composite (overlay) images are shown as applicable. Scale bars represent 10 μm. Images are representative of three independent replicates. (**b**) Quantification of Cy5 signal in biofilms. Relative abundance indicates the percentage of total biomass, as detected by brightfield, positive for Cy5 labeling. Values are normalized to *S. aureus* WT and shown as means ± S.E.M. from three independent replicates. No values were significantly different from WT by one-way ANOVA with Dunnett's post-test.

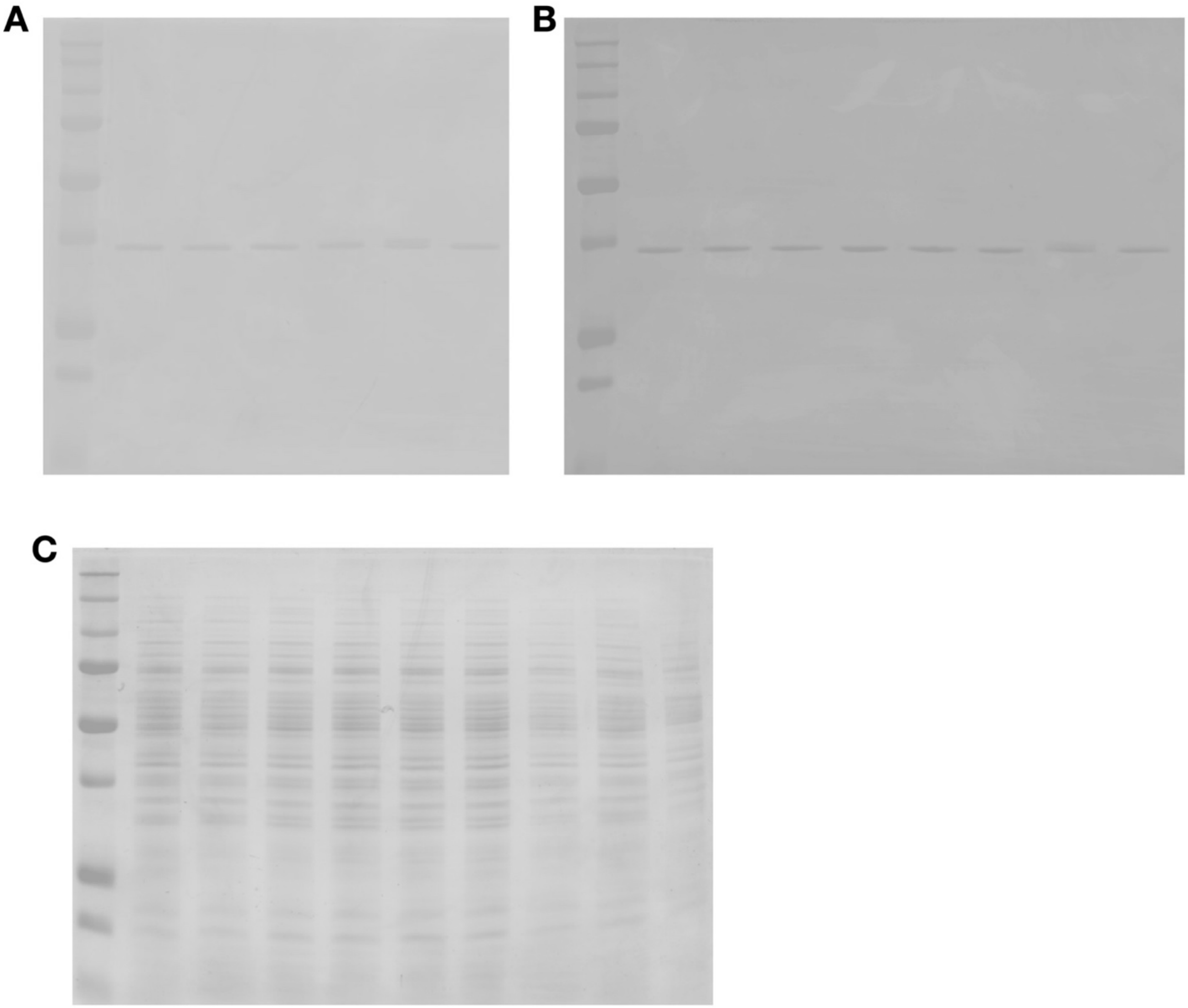


**Figure S7. Coomassie stains of SDS-PAGE gels.** (**a**) Corresponds to figure 1E. (**b**) Corresponds to figure S1. (**c**) Corresponds to figure 2A.

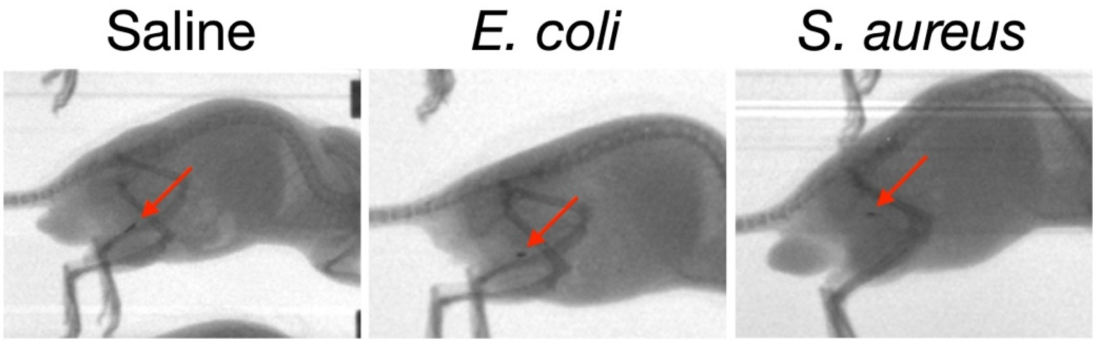


**Figure S8. CT scan of mice with titanium rods.** Computed tomography (CT) scan of representative mice to demonstrate localization of inserted titanium rods. The mice shown correspond to the mice shown in **Figure 4D**.

### Chemical synthesis

#### General methods

Unless otherwise noted, all reagents and solvents were purchased from commercial suppliers and used without further purification. Reactions were carried out under an argon atmosphere with continuous stirring unless specified otherwise. Reaction flasks were oven-dried at 100 °C overnight prior to use. Flash column chromatography was performed using SiliaFlash® P60 (230–400 mesh, SiliCycle) with reagent-grade solvents as indicated. Thin-layer chromatography (TLC) was conducted on pre-coated 0.25 mm silica gel plates (Merck) and spots were visualized under UV light. 1H and 13C NMR spectra were recorded on a BRUKER AVANCE III 500 spectrometer at room temperature (rt), using the indicated deuterated solvents. Chemical shifts (δ) are reported in parts per million (ppm) relative to tetramethylsilane (TMS) as an internal standard. 1H NMR data were reported in the order of chemical shift, multiplicity (s, singlet; brs, broad singlet; d, doublet; dd, doublet of doublets; t, triplet; td, triplet of doublets; q, quartet; qd, quartet of doublets; quint, quintet; m, multiplet and/or multiple resonance), number of protons, and coupling constant (J) in hertz (Hz). High-resolution mass spectra (HRMS) were analyzed by LC-ESI/MS on a Waters Acquity UPLC system coupled to a Thermo Exploris 240 Orbitrap mass spectrometer. The following abbreviations for reagents and solvents are used: N,N-dimethylformamide (DMF), trifluoroacetic acid (TFA).

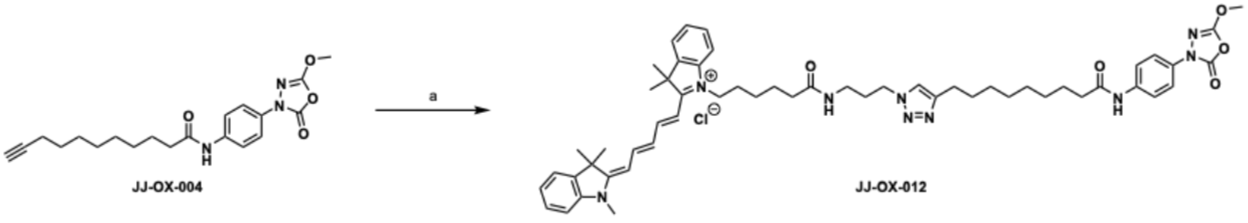


**Scheme S1.** Synthesis of **JJ-OX-012**. Reagent and conditions: (a) Cyanine5-N3, $CuSO_4 \cdot 5H_2O$, ascorbic acid, DMF, rt, 90%.

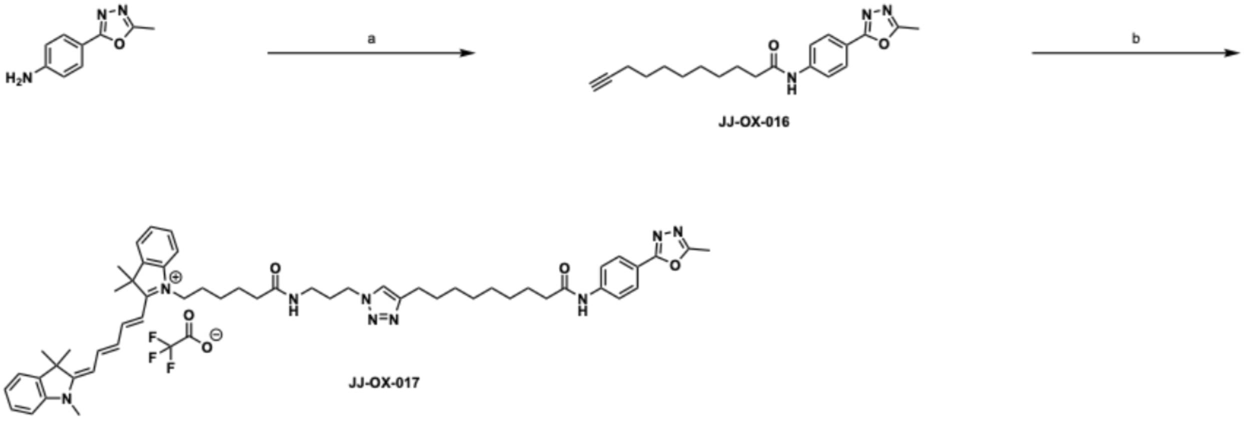

**Scheme S2.** Synthesis of **JJ-OX-016** and **JJ-OX-017**. Reagent and conditions: (a) 10-undecynoic acid, EDC·HCl, HOBt, DMF, 38%; (b) Cyanine5-N3, CuSO4·5H2O, ascorbic acid, DMF, rt, 14%.

**Synthesis of JJ-OX-012**

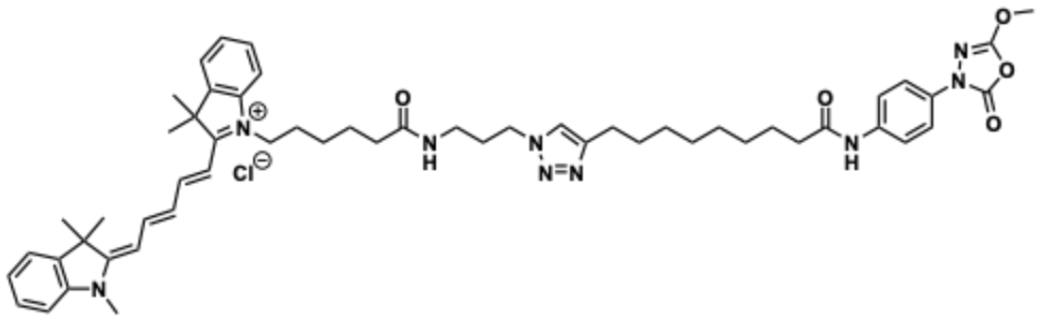

1-(6-((3-(4-(9-((4-(5-Methoxy-2-oxo-1,3,4-oxadiazol-3(2H)-yl)phenyl)amino)-9-oxononyl)-1H-1,2,3-triazol-1-yl)propyl)amino)-6-oxohexyl)-3,3-dimethyl-2-((1E,3E)-5-((E)-1,3,3-trimethylindolin-2-ylidene)penta-1,3-dien-1-yl)-3H-indol-1-ium chloride (**JJ-OX-012**)

To a stirred solution of **JJ-OX-004**[3] (7.5 mg, 20.0 µmol) in DMF (1.0 mL) was added cyanine5-azide (5.0 mg, 8.3 µmol) at ambient temperature. To a stirred reaction mixture, 100 mM CuSO4·5H2O solution in DMF (120 µL, 12.0 µmol) and 200 mM ascorbic acid solution in DMF (250 µL, 50.0 µmol) were added. After stirring at the same temperature for 32 h, the mixture was diluted with H2O, extracted 3 times with EtOAc, washed 3 times with brine. The combined organic layer was dried over Na2SO4 and concentrated in vacuo. The residue was purified by flash column chromatography on silica gel (MeOH/DCM = 1:10) to afford 7.2 mg (90%) of **JJ-OX-012** as a blue solid: 1H NMR (500 MHz, CDCl3) δ 9.68 (s, 1H), 8.53 (s, 1H), 7.99 (s, 1H), 7.90 – 7.87 (m, 2H), 7.79 – 7.71 (m, 2H), 7.65 – 7.62 (m, 2H), 7.40 – 7.36 (m, 1H), 7.34 – 7.32 (m, 1H), 7.27 – 7.14 (m, 4H), 7.09 (d, J = 7.9 Hz, 1H), 7.07 (d, J = 7.9 Hz, 1H), 6.68 (t, J = 12.5 Hz, 1H), 6.33 (d, J = 13.7 Hz, 1H), 6.15 (d, J = 13.5 Hz, 1H), 4.50 (t, J = 6.8 Hz, 2H), 4.07 (s, 3H), 3.99 (t, J = 7.6 Hz, 2H), 3.55 (s, 3H), 3.27 – 3.24 (m, 2H), 2.70 (t, J = 7.1 Hz, 2H), 2.51 – 2.48 (m, 2H), 2.41 (t, J = 7.3 Hz, 2H), 2.19 – 2.13 (m, 2H), 1.81 – 1.71 (m, 4H), 1.67 (s, 3H), 1.66 (s, 3H), 1.50 – 1.44 (m, 2H), 1.33 – 1.21 (m, 18H); HR-MS ($ESI^+$) *m/z*: $[M]^+$ calcd for $C_{55}H_{70}N_9O_5$ 936.5494; found 936.5476.

**Synthesis of JJ-OX-016 and JJ-OX-017**

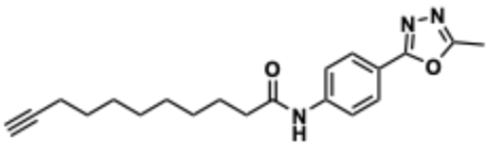

N-(4-(5-Methyl-1,3,4-oxadiazol-2-yl)phenyl)undec-10-ynamide (**JJ-OX-016**)

To a stirred solution of 4-(5-methyl-1,3,4-oxadiazol-2-yl)aniline (42.2 mg, 0.241 mmol) in DMF (3.00 mL) was added 10-undecynoic acid (43.9 mg, 0.241 mmol), EDC·HCl (51.0 mg, 0.266 mmol), and HOBt (40.7 mg, 0.266 mmol) at ambient temperature. After being stirred for 16 h at the same temperature, the reaction mixture was diluted with EtOAc and washed 3 times with brine. The organic layer was dried over Na2SO4 and concentrated in vacuo. The resulting residue was purified by flash column chromatography on silica gel (EtOAc/n-hexane = 2:1) to afford 31.3 mg (38%) of **JJ-OX-016** as a white solid: 1H NMR (500 MHz, CDCl3) δ 7.99 – 7.96 (m, 2H), 7.68 (d, J = 8.7 Hz, 2H), 7.36 (s, 1H), 2.61 (s, 3H), 2.39 (d, J = 7.6 Hz, 2H), 2.18 (td, J = 7.1, 2.7 Hz, 2H), 1.94 (t, J = 2.7 Hz, 1H), 1.74 (quint, J = 7.2 Hz, 2H), 1.52 (quint, J = 7.3 Hz, 2H), 1.43 – 1.30 (m, 8H); 13C NMR (126 MHz, CDCl3) δ 171.7, 164.7, 163.6, 141.1, 127.9, 119.7, 119.5, 84.9, 68.3, 38.0, 29.3, 29.3, 29.0, 28.8, 28.5, 25.6, 18.5, 11.3; HR-MS (ESI$^+$) *m*/*z*: [M + H]$^+$ calcd for $C_{20}H_{26}N_3O_2$ 340.2020; found 340.2017.

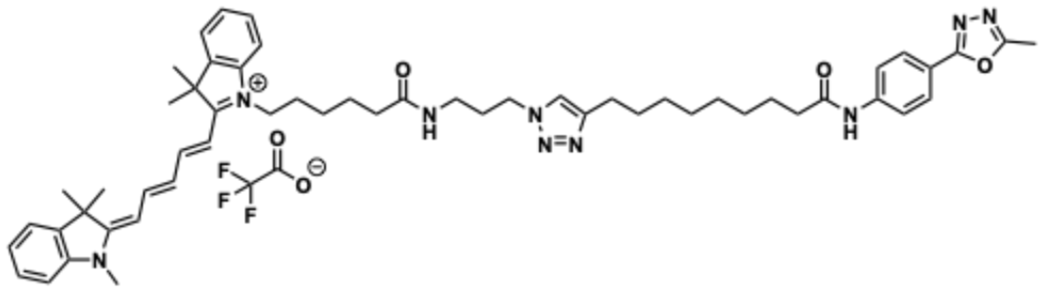


3,3-Dimethyl-1-(6-((3-(4-(9-((4-(5-methyl-1,3,4-oxadiazol-2-yl)phenyl)amino)-9-oxononyl)-1H-1,2,3-triazol-1-yl)propyl)amino)-6-oxohexyl)-2-((1E,3E)-5-((E)-1,3,3-trimethylindolin-2-ylidene)penta-1,3-dien-1-yl)-3H-indol-1-ium 2,2,2-trifluoroacetate (**JJ-OX-017**)

To a stirred solution of **JJ-OX-016** (6.8 mg, 20.0 μmol) in DMF (1.0 mL) was added cyanine5-azide (5.0 mg, 8.3 μmol) at ambient temperature. To a stirred reaction mixture, 100 mM CuSO4·5H2O solution in DMF (120 μL, 12.0 μmol) and 200 mM ascorbic acid solution in DMF (250 μL, 50.0 μmol) were added. After stirring at the same temperature for 32 h, the mixture was diluted with H2O, extracted 3 times with EtOAc, washed 3 times with brine. The combined organic layer was dried over Na2SO4 and concentrated in vacuo. The residue was filtered through a thin silica pad using MeOH, followed by purification with reverse-phase column chromatography (H2O/ACN = 95:5 to 5:95 gradient, 0.1% TFA) to afford 1.2 mg (14%) of **JJ-OX-017** as a blue solid: HR-MS (ESI$^+$) *m*/*z*: [M]$^+$ calcd for $C_{55}H_{70}N_9O_3$ 904.5596; found 904.5592.

### LC trace of JJ-OX-012

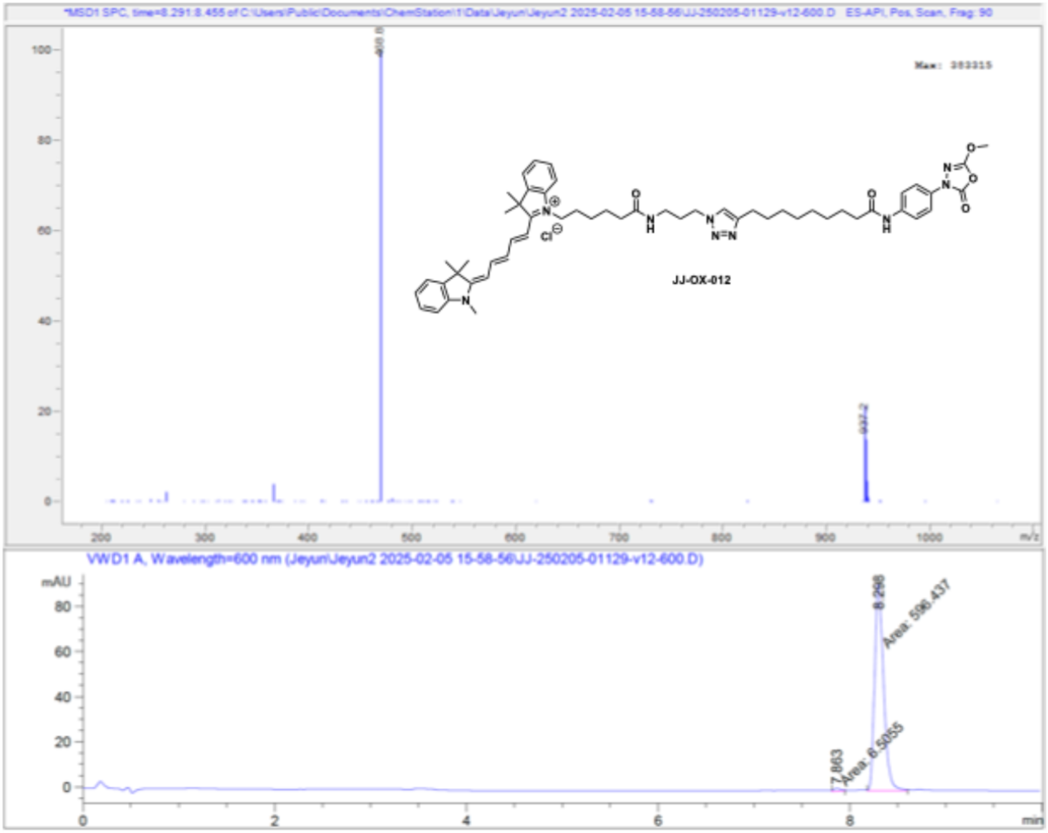


Signal 2: VWD1 A, Wavelength=600 nm

| Peak # | RetTime [min] | Type | Width [min] | Area [mAU*s] | Height [mAU] | Area % |
|---|---|---|---|---|---|---|
| 1 | 7.863 | MM T | 0.0896 | 6.50550 | 1.20969 | 1.0790 |
| 2 | 8.298 | MM T | 0.1087 | 596.43719 | 91.44978 | 98.9210 |
| Totals : | | | | 602.94270 | 92.65947 | |

### LC trace of JJ-OX-017

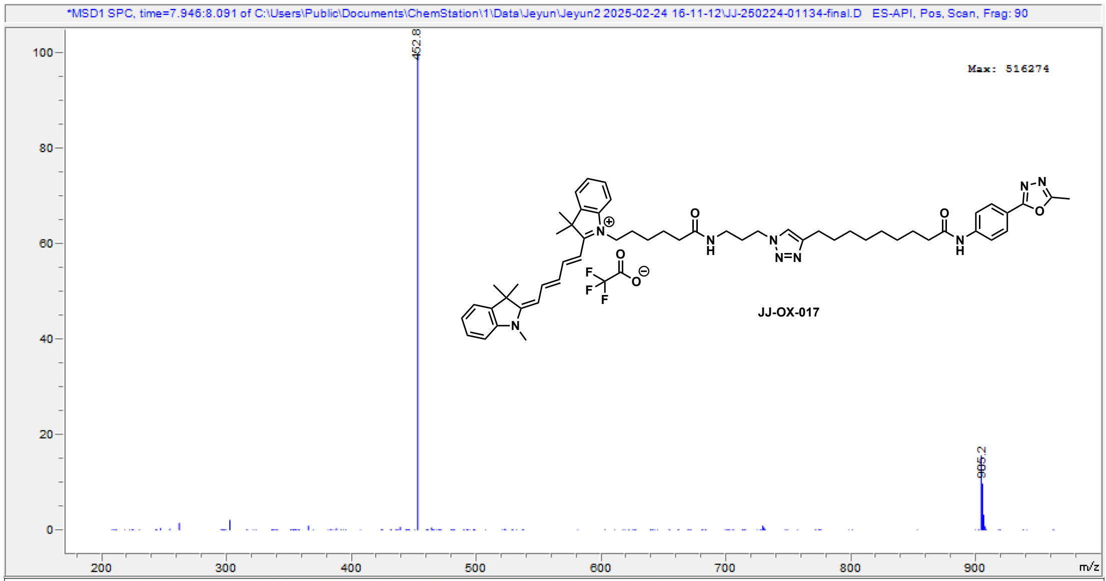


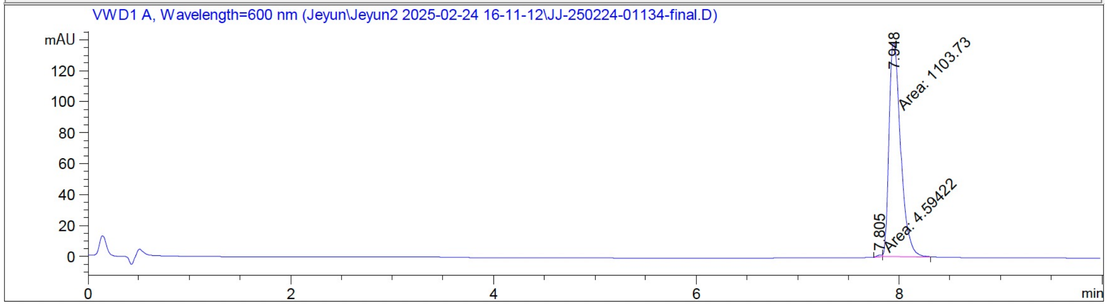


Signal 2: VWD1 A, Wavelength=600 nm

| Peak # | RetTime [min] | Type | Width [min] | Area [mAU*s] | Height [mAU] | Area % |
|---|---|---|---|---|---|---|
| 1 | 7.805 | MM T | 0.0644 | 4.59422 | 1.18821 | 0.4145 |
| 2 | 7.948 | MM T | 0.1328 | 1103.73401 | 138.53888 | 99.5855 |

Totals : 1108.32823 139.72709

**NMR Spectra**

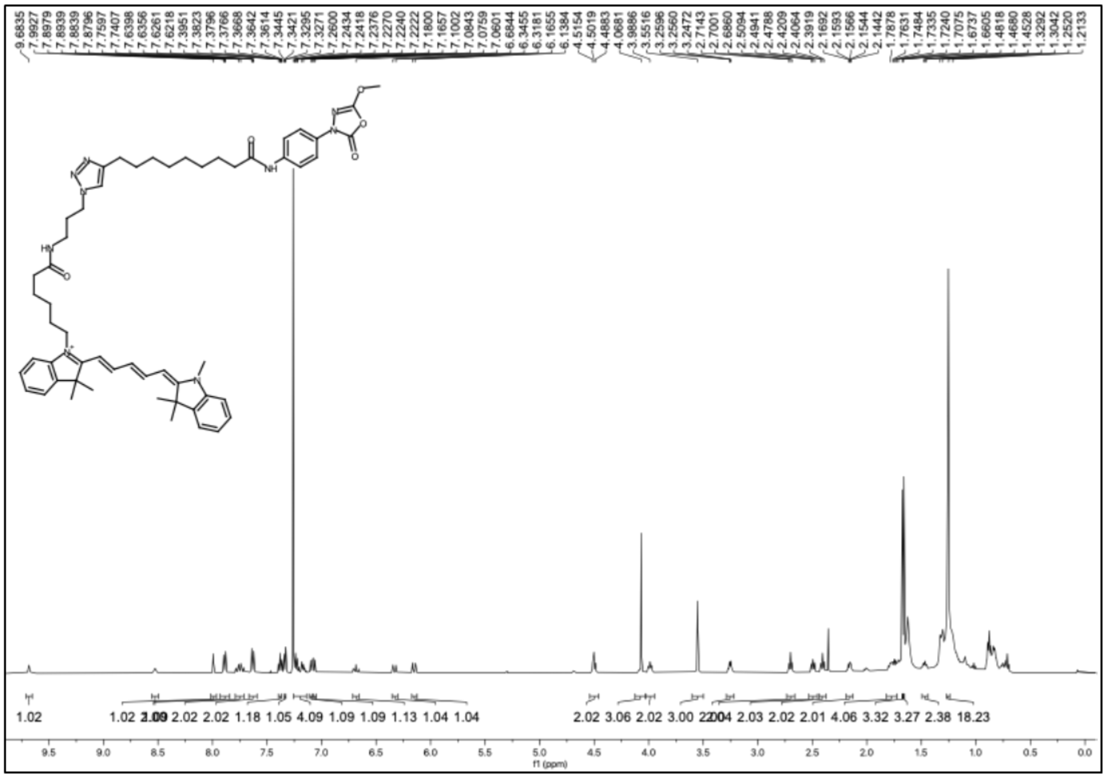

[1]H NMR spectrum of **JJ-OX-012** (CDCl3, 500 MHz)

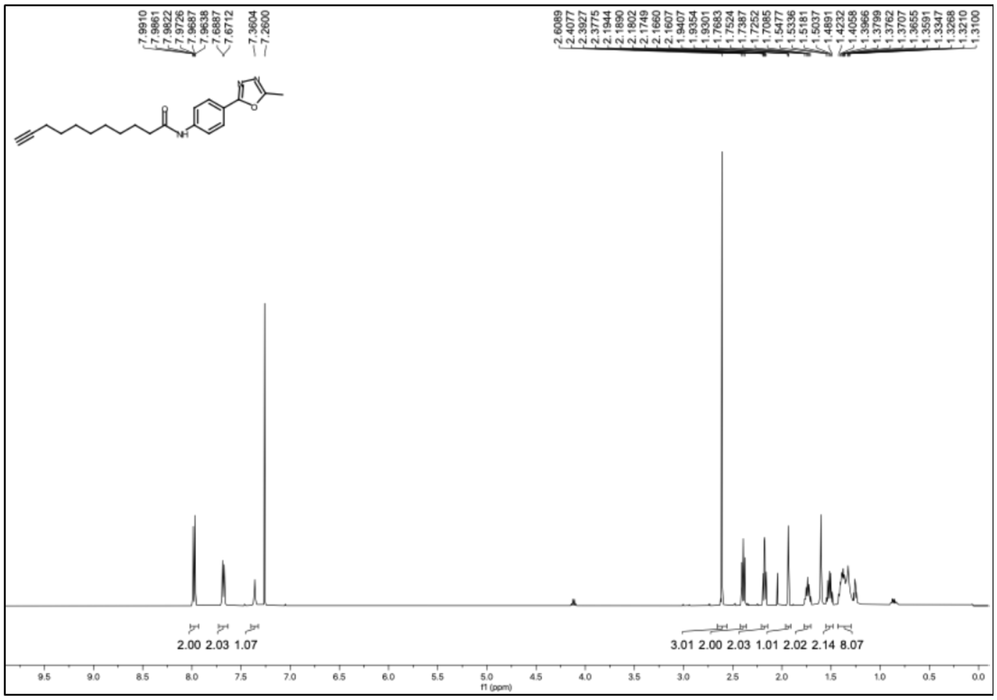

[1]H NMR spectrum of **JJ-OX-016** (CDCl3, 500 MHz)

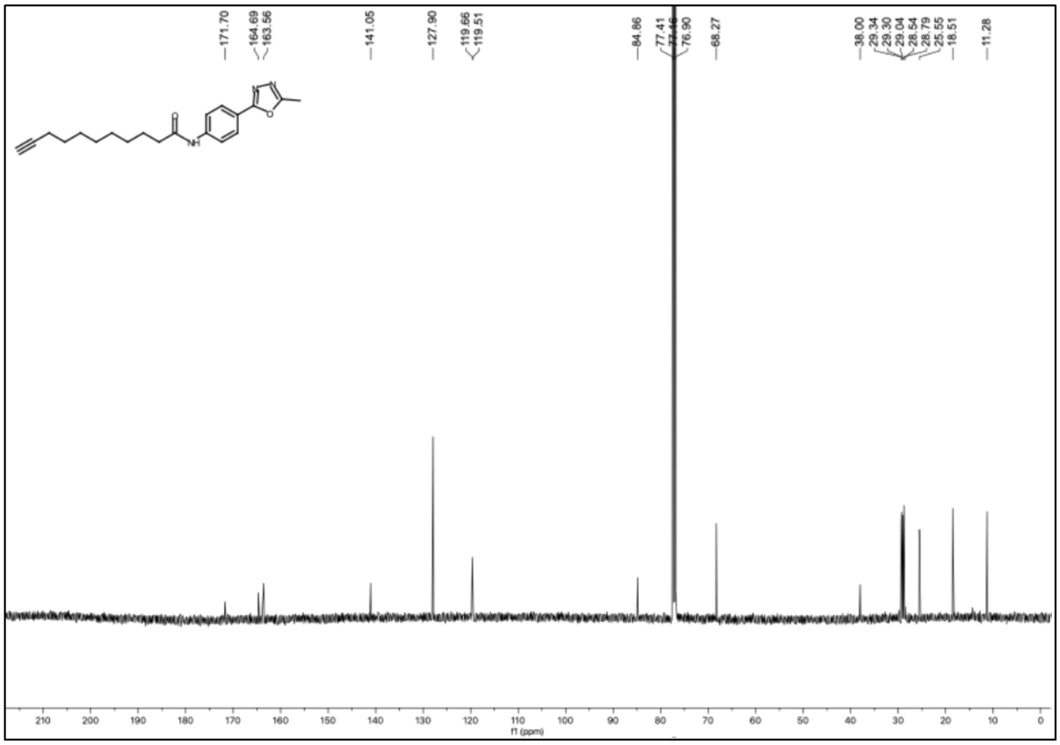


[13]C NMR spectrum of **JJ-OX-016** (CDCl3, 126 MHz)
